# Energetic and Environmental Constraints on the Community Structure of Benthic Microbial Mats in Lake Fryxell, Antarctica

**DOI:** 10.1101/687103

**Authors:** Megan L. Dillon, Ian Hawes, Anne D. Jungblut, Tyler J. Mackey, Jonathan A. Eisen, Peter T. Doran, Dawn Y. Sumner

**Affiliations:** Lawrence Berkeley National Laboratory Climate and Ecosystem Sciences Division 70A-2245B, One Cyclotron Rd Berkeley, CA 94720; Department of Earth and Planetary Sciences, University of California, Davis; Coastal Marine Field Station, University of Waikato; Life Sciences Department, Natural History Museum; Department of Earth, Atmospheric, and Planetary Sciences, Massachusetts Institute of Technology; Department of Evolution and Ecology, University of California, Davis; Geology and Geophysics, Louisiana State University

## Abstract

Ecological communities are commonly thought to be controlled by the dynamics of energy flow through environments. Two of the most important energetic constraints on all communities are photosynthetically active radiation (PAR) and oxygen concentration ([O_2_]). Microbial mats growing on the bottom of Lake Fryxell, Antarctica, span environmental gradients in PAR and [O_2_], which we used to test the extent to which each controls community structure. Metagenomic analyses showed variation in the diversity and relative abundances of Archaea, Bacteria, and Eukaryotes across three [O_2_] and PAR conditions. Where [O_2_] saturated the mats or was absent from the overlying water, PAR structured the community. Where [O_2_] varied within mats, microbial communities changed across covarying PAR and [O_2_] gradients. Diversity negatively correlated with [O_2_] and PAR through mat layers in each habitat suggesting that, on the millimeter-scale, communities are structured by the optimization of energy use. In contrast, [O_2_] positively correlated with diversity and affected the distribution of dominant populations across the three habitats, suggesting that meter-scale diversity is structured by energy availability. The benthic microbial communities in Lake Fryxell are thus structured by energy flow in a scale-dependent manner.

## Introduction

The composition and diversity of biological communities respond strongly to energy availability, as evidenced by the positive correlation between net primary productivity (NPP) and diversity (Gillman *et al.* 2015). This correlation led to the development of species-energy theory, which posits that habitats with greater energy capture by a community have more niche diversity (Hurlbert and Stegen 2014). The relationships between communities and environmental characteristics have historically been studied in the presence of macroflora and -fauna because there are few places on Earth where microbial niches are not at least partly structured by macroorganisms. Some of these environments include hot springs (Amin *et al.* 2017), the deep biosphere (Ino *et al.* 2018), and snowfall (Brown and Jumpponen 2019), and studies of them have been foundational to evaluating the application of ecological principles to microbial ecosystems.

Species-energy theory is likely applicable to microbial ecosystems based on recent experimental work investigating the effects of increased carbon substrate availability on community composition and diversity in a groundwater system (Zhou *et al.* 2014). Other laboratory and field studies also suggest that microbial diversity increases with energy availability (Bernstein *et al.* 2017) and [O_2_] (Spietz *et al.* 2015). However, the actual energy available to a community is a complex function of both the biogeochemistry of the environment (*e.g.*, (Marlow *et al.* 2014)) and the metabolisms that can be utilized by the microbes present (Tilman 1994). When and where specific groups of microbes are highly efficient at capturing available energy, the effects of energy flow on microbial diversity are complex. One important source of energy for many microbial ecosystems is photosynthetically active radiation (PAR). Where PAR is high, species-energy theory predicts that ecosystems will have increased taxonomic richness and *vice versa*. For example, where PAR is available, oxygenic photosynthesizers are particularly abundant, and should drive NPP in their habitats. These primary producers simultaneously affect the redox state of electron donors and acceptors (Macalady *et al.* 2013), thus increasing resource availability for heterotrophic community members. However, ecosystems with a large population of oxygenic photosynthesizers sometimes show negative correlations between alpha diversity and primary productivity (13). In these communities, oxygenic photosynthesizers capture the majority of the energy in a habitat resulting in greater population sizes of oxygenic photosynthesizers and reduced relative abundance of heterotrophs, anaerobes, and overall alpha diversity, in conflict with the species-energy theory (Hurlbert and Stegen 2014).

An alternative to the species-energy theory - the maximum power principle - argues that communities may be structured to optimize productivity (Odum and Pinkerton 1955; DeLong 2008). The maximum power principle posits that communities that maximize the capture of energy will be at a selective advantage over those that do not, and the relative abundances of populations in an ecosystem will adjust accordingly. In contrast to species-energy arguments, the maximum power principle predicts that optimisation of energy capture may result in decreased diversity (Odum and Pinkerton 1955; DeLong 2008). Both species-energy theory and the maximum power principle assert that ecological communities are structured by resource heterogeneity (spatial, temporal, or both) and both hypotheses are therefore variations of the species sorting model according to metacommunity theory (Leibold and Chase 2017).

Understanding relationships between energy flow and diversity in microbial ecosystems is crucial to distinguishing what structures them. The ecological theories discussed above were developed from well-studied macroscopic ecosystems. If they are universal, they should apply to microbial ecosystems. By tracking microbial diversity across gradients in total and type of energy available, we can gain insights into the similarities and differences in how ecosystems are structured without macroscopic organisms. Specifically, we can use such gradients to determine whether the diversity of microbial ecosystems is structured by, and correlated with, energy availability and harvesting. However, we can expect the composition and diversity of microbial communities to also be constrained by the biogeochemistry of the local environment, given the diversity of metabolic strategies employed by microorganisms. Thus, heterogeneity of resources may be more important than total energy input in determining alpha diversity of a microbial ecosystem.

The processes determining the structure of microbial ecosystems have been investigated largely in extreme environments such as those with high salinity (Kunin *et al.* 2008; Schneider *et al.* 2013) or high temperatures (Vick *et al.* 2010; Schneider *et al.* 2013; Fortney *et al.* 2016), where macroorganisms are excluded and microorganisms predominate. Similarly, ice-covered lakes in Antarctica are extensive habitats for microbial mats that grow in the absence of most plants and animals (Van Trappen *et al.* 2002; Taton *et al.* 2003; Rios *et al.* 2004). The circumpolar Southern Ocean currents and the freezing temperatures of the continent isolate Antarctica from much of the terrestrial plant and animal life in the rest of the world (20), and the extreme aridity of the McMurdo Dry Valleys (MDVs) further limits the diversity of organisms that might otherwise inhabit the lakes (Cary *et al.* 2010).

McMurdo Dry Valley Lake Fryxell is a perennially ice-covered, density-stratified lake with a steep oxycline and gently sloping basin (Lawrence and Hendy 1985). Steep PAR and [O_2_] gradients existed from 9 to 10 m depth in the lake in 2012 (Figure 1). The persistent density gradients in the water column lead to stratification in Lake Fryxell’s pelagic community (Spaulding *et al.* 1994): NPP in the photic zone is dominated by oxygenic photosynthesis, but the redox structure of the water column and the distribution of irradiance allows anoxygenic photosynthesis to contribute to primary productivity as well (Klatt *et al.* 2015). The distribution of pelagic chemolithoautotrophs in Lake Fryxell is also limited to depths greater than 11 m by redox potential (Dolhi *et al.* 2015).

**Figure 1.**
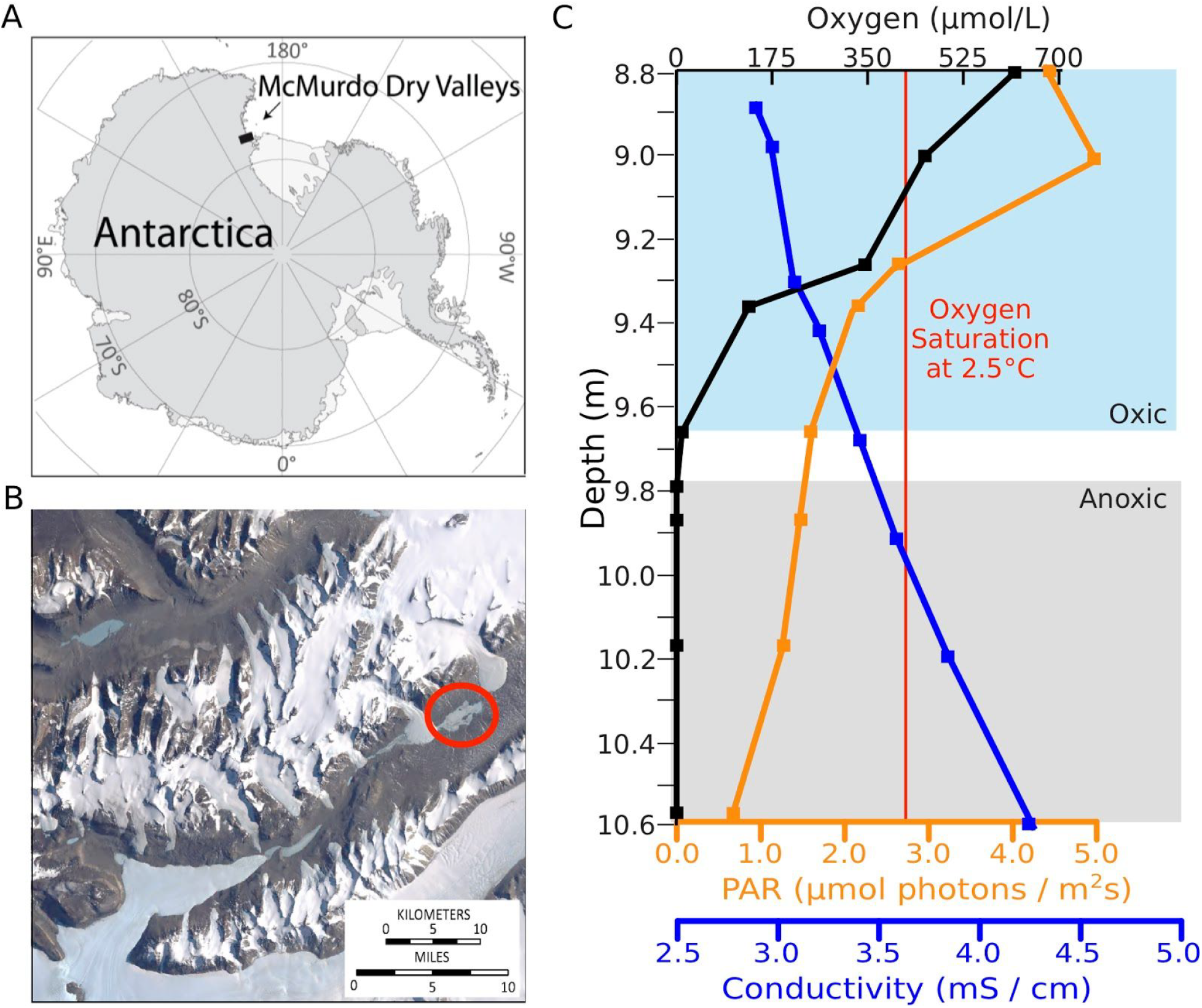
A) The McMurdo Dry Valleys are in Southern Victorialand, East Antarctica. B) Location of Lake Fryxell in Taylor Valley (circled in red) (Herried B. 2010). C) Oxygen concentration and conductivity, PAR, and oxygen saturation at 0°C along a benthic mat transect in Lake Fryxell in November 2012 (Jungblut *et al.* 2016).

Together, the irradiance and oxygen gradients in Lake Fryxell, and the stratification of the pelagic community, indicate that PAR and [O_2_] are key environmental characteristics that influence the community composition and diversity. Lake Fryxell is particularly informative, as the distinct gradients in PAR and [O_2_] do not co-vary across all habitats, allowing for comparison of these parameters. The variation of PAR and [O_2_] in Lake Fryxell provides an opportunity to study how the structure of benthic microbial mat communities differ according to habitat, specifically energy availability. In this study, we present the benthic community diversity and composition in Lake Fryxell with the aim of providing insights into the relative importance of PAR and [O_2_] in shaping community structure and testing ecological theories of energy use in a microbial ecosystem.

## Methods

### Site Description

Lake Fryxell (77° 36’S 162° 6’ E) is a density stratified and oligotrophic freshwater ecosystem in the MDVs, Antarctica. It is 5 x 1.5 km in extent and the maximum depth is approximately 20 m (Green and Lyons 2009). Water is supplied to Lake Fryxell by 13 glacial melt-water streams, with most water coming from the Canada and Commonwealth glaciers (McKnight *et al.* 1999). Water is lost by evaporation and ablation from the surface, and changes in lake level reflect net water balance; there are no out-flowing streams (Lawrence and Hendy 1985).

Environmental conditions in Lake Fryxell are strongly affected by a 4-5 m thick perennial ice cover (Obryk *et al.* 2016). The ice cover inhibits wind mixing and prevents gas equilibration between lake water and the atmosphere. A lack of vertical mixing is due to a stable density stratification, as demonstrated by conductivity profiles (Priscu 2014; Jungblut *et al.* 2016) (Figure 1). Stratification limits the transport of nutrients and redox pairs to diffusion and creates stable redox and nutrient gradients in the water column (Lee *et al.* 2004; Jungblut *et al.* 2016).

Oxygen and other gases are excluded as ice freezes onto the underside of the ice cover during winter, building to gas supersaturation in shallow waters (Hood, Howes and Jenkins 1998). Oxygen concentration declines rapidly with depth below 8 m, and oxygen is absent from the water column below approximately 9.7 m (Sumner *et al.* 2015; Jungblut *et al.* 2016). In addition to contributing to gas buildup in shallow waters, the ice cover also limits the amount of light reaching the lake waters. During the summer, the ice cover transmits approximately 1 % of incident irradiance (30), which provides the lake’s primary energy influx. Light reaching the floor of Lake Fryxell declines with increasing depth in the water column but is adequate to support photosynthesis at the top of benthic mats to depths of 11 m during the summer months (Sumner *et al.* 2015; Jungblut *et al.* 2016). Temperature varies from 2.4 - 2.7°C and pH varies from 7.44 - 7.52 from 8.9 to 11.0 m depth in the lake along an established transect (Hillman 2013; Jungblut *et al.* 2016). All depths are in relation to lake level during November 2012.

Lake Fryxell supports a rich microbial community. The pelagic community thrives near the deep chlorophyll maximum, located just above the oxic-anoxic transition (9-10 m), which is localized where nutrients diffuse up into the photic zone (Roberts 2017). Benthic microbial mats grow to depths of at least 10.5 m, forming cm-to-dm-scale thick mats (Jungblut *et al.* 2016). Mat pigmentation and morphology transition from purple pinnacles to orange ridges and pits to bright green flat mats with depth, as PAR and [O_2_] decline (Jungblut *et al.* 2016). Oxygenic photosynthesis raises redox conditions seasonally within mats near the oxic-anoxic transition in the water column (Sumner *et al.* 2015). Microelectrode profiles taken in late spring through the benthic microbial mats showed that [O_2_] varied at different water depths (Sumner *et al.* 2015). At 9.0 m, [O_2_] ranged from ~650 μmol O_2_/ L in the water immediately above and just below the surface of the mat, to ~825 μmol O_2_/ L at 2 mm below the mat surface. The mats still contained > 500 µmol O_2_/ L at 17 mm below the surface. In contrast, at 9.8 m, there were 0 μmol O_2_/ L in the overlying water column, but up to 50 μmol O_2_/ L were present within the upper mat due to photosynthetic O_2_ production, falling back to 0 μmol O_2_/ L by 6 mm depth into the mat (Sumner *et al.* 2015).

### Sampling

The benthic microbial mats in Lake Fryxell, Antarctica were sampled in November 2012, as described by ref. 30. Sampling was performed at 9.0, 9.3, and 9.8 m depths along a transect that was installed in 2006 (Hillman 2013). Divers retrieved samples from the bottom of the lake by cutting samples out of *in situ* mats using a spatula and lifting them into plastic boxes underwater. Upon delivery to the surface, multiple biological replicates of samples from each depth were dissected according to layer pigmentation and morphology using sterile sampling equipment, resulting in three layers at 9.0 m (top, middle, bottom), three layers at 9.3 m, and four layers at 9.8 m (film, top, middle, bottom). Mats at 9.3 m had complicated topography with 0.5-1.0 cm deep pits between ridges. At this depth, samples were collected from the tops of the ridges (top), within the ridges including their edges (middle), and from the bottoms of pits (bottom) ((Jungblut *et al.* 2016) Supplementary Information). The samples were preserved in the field immediately after sampling using an Xpedition Soil/Fecal DNA MiniPrep kit (Zymo Research, Irvine, CA) frozen, and shipped frozen to UC Davis where they were stored at −80 °C until DNA was extracted.

### Metagenomic Sequencing

DNA was extracted using an Xpedition Soil/Fecal DNA MiniPrep kit (Zymo Research, Irvine, CA) as per manufacturer instructions from biological and technical replicates of 10 sample types (Table S1). Metagenomic sequencing was performed at the University of California, Davis Genome Center DNA Technologies Core (http://dnatech.genomecenter.ucdavis.edu/) using the Illumina HiSeq 2500, PE 250 platform. Library preparation was performed using Illumina’s Nextera DNA Kit (Oligonucleotide sequences © 2007-2013 Illumina, Inc.). Reads were quality filtered to Q20 using FASTX-Toolkit v0.0.13 (Gordon, Hannon and Others 2010), and forward and reverse reads were joined using PEAR v0.9.6 (Zhang *et al.* 2014). Fastq files with fewer than 10000 reads were determined to be outliers and were therefore removed from downstream analyses. Metagenomic reads are available via NCBI’s sequence read archive (PRJNA291280).

### Taxonomic Assignment

Reads were assigned to Bacteria, Archaea, and Eukaryotes. Protein-level classifications for taxonomic assignments were made per replicate using Kaiju v1.4.5 (Menzel, Ng and Krogh 2016) and NCBI’s non-redundant protein database (accessed 3/2/2017), including Bacteria, Archaea, Eukaryotes, and Viruses. All assignments, from domain to operational taxonomic units (OTUs), were made using Kaiju’s greedy mode with 5 allowed substitutions, a minimum match score of 70, and low-complexity query sequence filtering. Taxa in relative abundance less than 0.5 % were removed from downstream analyses.

### Phylogenetic Diversity Analyses

Phylosift v1.0.1 (Darling *et al.* 2014) was used to generate phylogenetic placements for the marker genes (https://phylosift.wordpress.com/tutorials/scripts-markers/) per replicate using default settings. Guppy v1.1 was used to calculate balance weighted phylogenetic diversity (bwpd, alpha diversity) (Matsen, Kodner and Virginia Armbrust 2010), which provides alpha diversity based on phylogenetic relationships using .jplace files generated by Phylosift as input, as opposed to community count data (Nipperess and Matsen 2013). Guppy v1.1 was also used to perform edge principal components analysis (EPCA, beta diversity) treating every pquery as a point mass concentrated on the highest-weight placement and rotating three dimensions for support overlap minimization. Unrooted phylogenetic diversity rarefaction curves were generated using Guppy (McCoy and Matsen 2013).

### Statistical Analyses

Significant differences in taxonomic classification between samples were determined using Permutational Multiple Analysis of Variance (PERMANOVA) in R v3.3.2 (Warton, Wright and Wang 2012; Anderson and Walsh 2013; R Core Team and Others 2013). Lineages determined to differ significantly between samples were then subjected to Analysis of Variance and Tukey’s Honest Significant Difference (Tukey’s HSD) test (Tukey 1949) in R v3.3.2 (R Core Team and Others 2013) to establish which lineages differed among samples.

## Results

### Taxonomic Annotation

After quality filtering, the total amount of metagenomic sequence data came to approximately 47.1 billion bp from approximately 177 million reads (Table S1). Taxa from Bacteria, Archaea, and Eukarya were present in all samples (Figure 2, Tables S3–5). The most relatively abundant organisms across all samples are from the domain bacteria and the phyla Proteobacteria (19.2%) and Cyanobacteria (7.5%) (Figure 3). The most abundant genera across all samples belong to the phylum Cyanobacteria, and genera in Cyanobacterial subsection III, filamentous Cyanobacteria that lack cell differentiation (Tomitani *et al.* 2006), are dominant. One genus is found in far greater abundance (P < 0.01, Table S5) than any other taxon in the film, top, and middle layers of the microbial mat growing at 9.8 m, identified as a species of *Phormidium* in previous 16S rRNA gene analyses (ref. 30 Supplementary Information); here, we also refer to this genus as *Phormidium*, even though reads in this genus were annotated as both *Phormidium* and *Oscillatoria* (Altschul *et al.* 1990). In this study, 55 *Phormidium* OTUs occur across all ten sample types, but *Phormidium pseudopristleyi* is found in far greater relative abundance in the film at 9.8 m than in any other sample (Table S5).

**Figure 2.**
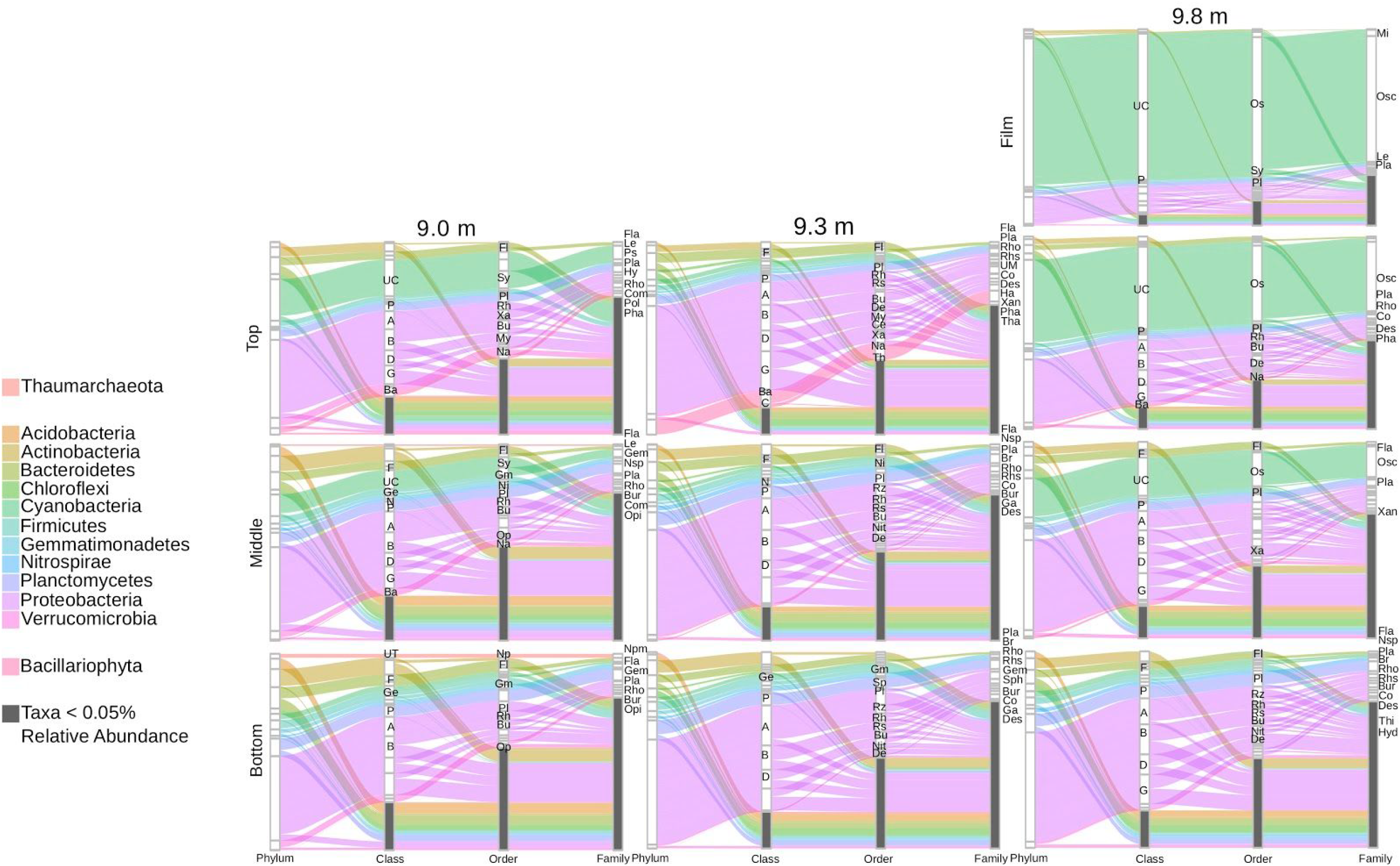
Relative abundance of taxa from all samples. The relative proportion of Cyanobacteria to Proteobacteria decrease through mat layers at all depths and mats at 9.8 m are dominated by the family Oscillatoriaceae. Taxa labels: A, Alphaproteobacteria; B, Betaproteobacteria; Ba, Bacillariophyceae; C, Coscinodiscophyceae; D, Deltaproteobacteria; F Flavobacteriia; G, Gammaproteobacteria; Ge, Gemmatimonadetes; N, Nitrospirales; P, Planctomycetica; UC, Undefined Cyanobacteria Class; UT, Undefined Thaumarchaeota Class; Bu, Burkholderiales; Ce, Cellvibrionales; De, Desulfobacterales; Fl, Flavobacteriales; Gm, Gemmatimonadales; My, Myxococcales; Na, Naviculales; Ni, Nitrospirales; Np, Nitrospumilales; Nit, Nitrosomonadales; Os, Oscillatoriales; Op, Optitutales; Pl, Planctomycetales; Rh, Rhodobacterales; Rs, Rhodospirilalles; Rz, Rhizobiales; Sp, Sphingomonadales; Sy, Synechococcales; Th, Thalassiosirales; Xa, Xanthomonadales; Br, Bradyrhizobiaceae; Bur, Burkholderiaceae; Co, Comamonadaceae; Des, Desulfobacteraceae; Fla, Flavobacteriaceae; Ga, Gallionellaceae; Gem, Gemmatimonadaceae; Hy, Hyphomonadaceae; Hyd, Hydrogenophilaceae; Le, Leptolyngbyaceae; Mi, Microcoleaceae; Nsp, Nitrospiraceae; Npm, Nitrosopumilaceae; Osc, Oscillatoriaceae; Opi, Opitutaceae; Pha, Phaeodactylaceae; Po, Polyangiaceae; Pla, Planctomycetaceae; Ps, Pseudanabenaceae; Rho, Rhodobacteraceae; Rhs, Rhodospirilaceae; Sph, Sphingomonadaceae; Tha, Thalassiosiraceae; Thi, Thiobacillaceae; Xan, Xanthomonadaceae; UM, Undefined Myxococcales Family.

**Figure 3.**
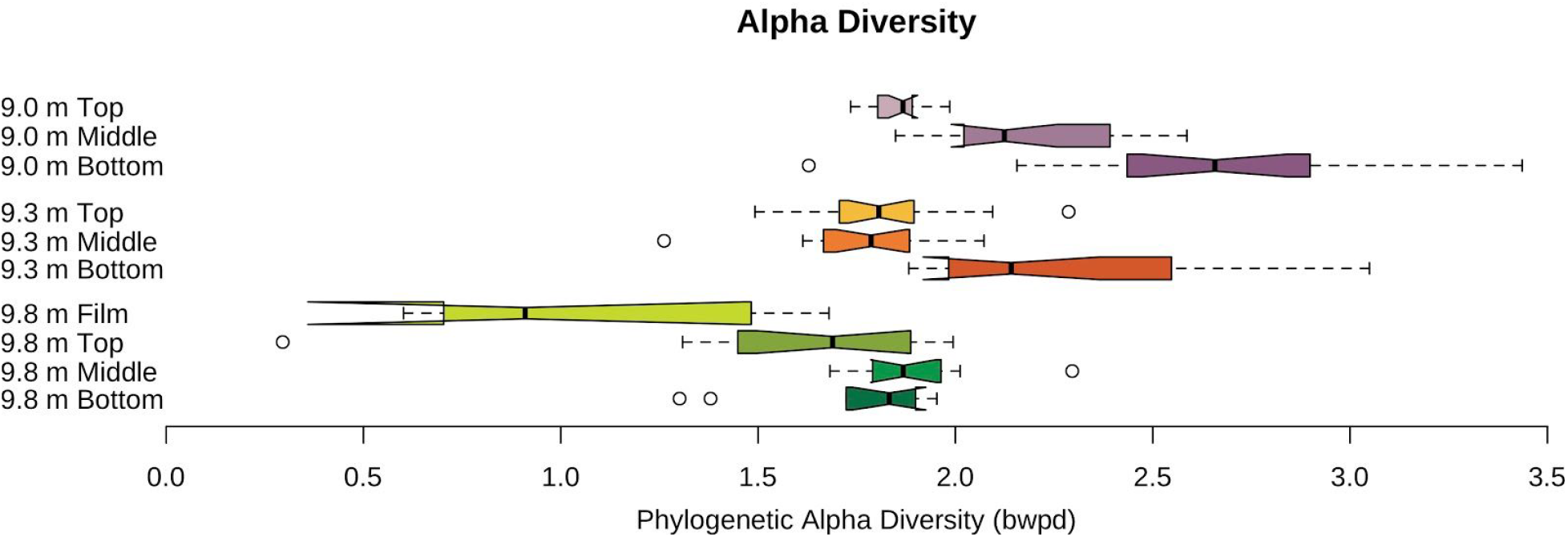
Alpha diversity of microbial mat communities by lake depth and mat layer as measured by balance-weighted phylogenetic diversity, which provides alpha diversity based on phylogenetic relationships as opposed to community count data. Whiskers represent extreme values, and boxes represent the interquartile range.

### Alpha Diversity

Alpha diversity varies both with distance into the mat and with water depth (Figure 3, Table 1). Alpha diversity decreases with depth in the lake when layers are grouped at a specific depth (Figure 3), positively correlating with both PAR and [O_2_]. In contrast, alpha diversity increases into the mat from the top to bottom layers when depths are pooled (Figure 3), negatively correlating with both PAR and [O_2_]. In 9.0 m mats, where oxygen is supersaturated and effectively constant throughout the mat and PAR decreases through the mat, alpha diversity increases with depth (Figure 3). The diversity of the dark bottom layers from mats at each depth differ by [O_2_]. At 9.0 m, where [O_2_] is highest, the alpha diversity of the bottom layer is highest and at 9.8 m, where [O_2_] is lowest, the alpha diversity of the bottom layer is lowest (Figure 3). Simpson’s diversity indices does not vary significantly between sample types (Table S2).

**Table 1.** Adjusted P-values of alpha diversity (bwpd, calculated using abundance-weighted phylogenetic placements) comparisons between samples, calculated using Tukey’s HSD. Mat layer communities were considered significantly different at P-values < 0.01. Comparisons without significant differences are marked -, and self-comparisons are marked x.

|  | 9.0 m<br>Top | 9.0 m<br>Middle | 9.0 m<br>Bottom | 9.3 m<br>Top | 9.3 m<br>Middle | 9.3 m<br>Bottom | 9.8 m<br>Film | 9.8 m<br>Top | 9.8 m<br>Middle | 9.8 m<br>Bottom |
| --- | --- | --- | --- | --- | --- | --- | --- | --- | --- | --- |
| 9.0 m<br>Top | x |  |  |  |  |  |  |  |  |  |
| 9.0 m<br>Middle | - | x |  |  |  |  |  |  |  |  |
| 9.0 m<br>Bottom | 0.00E+00 | 1.85E-03 | x |  |  |  |  |  |  |  |
| 9.3 m<br>Top | - | - | 0.00E+00 | x |  |  |  |  |  |  |
| 9.3 m<br>Middle | - | 5.10E-03 | 0.00E+00 | - | x |  |  |  |  |  |
| 9.3 m<br>Bottom | - | - | - | 9.54E-03 | 1.96E-03 | x |  |  |  |  |
| 9.8 m<br>Film | 1.66E-04 | 0.00E+00 | 0.00E+00 | 4.00E-04 | 1.34E-03 | 0.00E+00 | x |  |  |  |
| 9.8 m<br>Top | - | 7.80E-06 | 0.00E+00 | - | - | 3.10E-06 | - | x |  |  |
| 9.8 m<br>Middle | - | - | 1.00E-07 | - | - | - | 1.34E-04 | - | x |  |
| 9.8 m<br>Bottom | - | - | 0.00E+00 | - | - | 7.33E-03 | - | - | - | x |

### Beta Diversity

Tukey’s HSD test and PERMANOVA revealed the OTUs contributing to statistically significant differences between samples (Tables S3–5). At 9.0 m, 54 OTUs (approximately 6% of the OTUs identified throughout the 9.0 m mats) vary significantly (P < 0.01) between layers (Table S3, Figure 2), including several bacterial OTUs as well as diatoms and one archaeal genus (*Nitrosopumilis*). At 9.3 m, only eight bacterial and archaeal OTUs (0.1% of the OTUs identified throughout the 9.3 m mats) vary significantly (P < 0.01) between the layers (Table S4, Figure 2). At 9.8 m, one OTU (0.5% of the OTUs identified throughout the 9.8 m mats), *Phormidium pseudopristleyi*, vary significantly (P < 0.01) between the layers (Table S5, Figure 2).

Edge Principal Components Analysis (EPCA) illustrates the differences in community composition among samples (Figure 4). At 9.0 and 9.3 m, samples from the same mat layer group together (Figure 4B, Figure 4C). At 9.8 m, the samples do not group by top, middle or bottom layers (Figure 4D).

**Figure 4.**
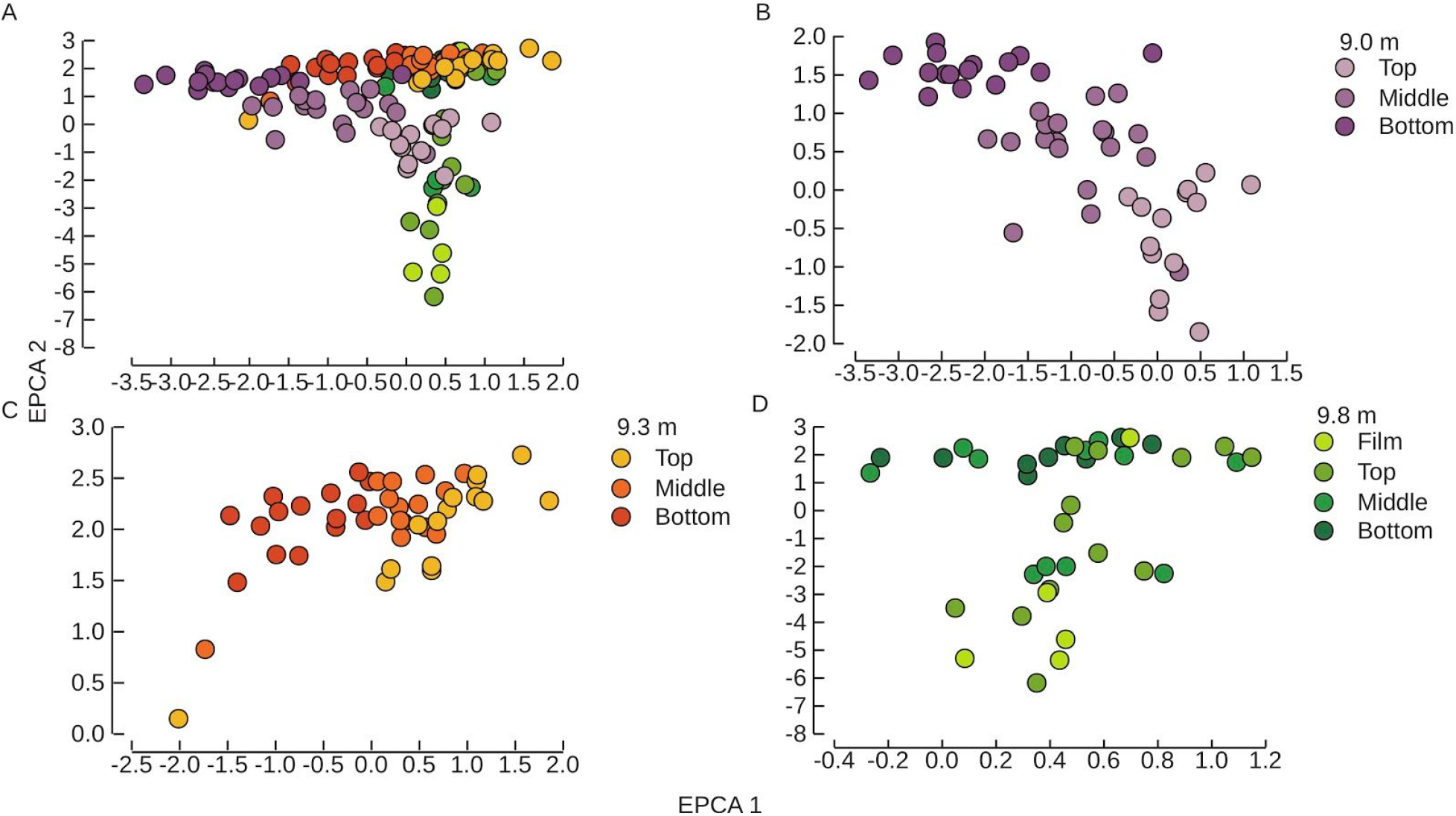
Beta diversity analysis using EPCA of mat layer communities by depth as tested by PERMANOVA and Tukey’s HSD (Tukey, 1949) (Tables 2-4). (A) All samples, (B) samples from all mat layers at 9.0 m (C) samples from all mat layers at 9.3 m, and (D) samples from all mat layers at 9.8 m.

In summary, oxygen-supersaturated mats (9.0 m) have higher alpha diversity (approximately 1.7 – 2.6, Figure 3) and larger phylogenetic differences in community composition between layers (Figure 4), whereas anoxic mats (9.8 m) have lower alpha diversity (approximately 0.8 – 1.9, Figure 3) and less phylogenetic differences in community composition between layers (Figure 4). In contrast, top layers have fewer taxonomic differences across lake depths than subsurface layers (Figure 2).

## Discussion

### Effects of [O_2_] and PAR on community composition and diversity

In Lake Fryxell, PAR and [O_2_] influence the microbial community in the various layers of the benthic mats differently, depending on the lake depth. The data suggest that the 9.0 m mats were structured by PAR. All layers grew at supersaturated [O_2_] (Sumner *et al.* 2015), and the mat communities from the 9.0 m habitat likely did not experience physiologically relevant variation in [O_2_]. The decrease in predicted photoautotrophic Cyanobacteria through the layers at 9.0 m (Figure 3) can be explained by the top layer of the mat being illuminated and underlying layers being shaded. Thus, differences in community composition (Figure 4) and diversity (Figure 3) from the top to the bottom samples at 9.0 m can be attributed to a decline in PAR with depth into the mat.

The mats at 9.3 m had complex topography with pits and ridges, and all sample types (top, middle, and bottom) included some mat exposed to lake water ((Jungblut *et al.* 2016) Supplementary Figure S2B). The mat topography likely affected PAR and [O_2_] exposure. Although PAR was not measured in the pits, some light penetrated into them, based on downward-looking photographs in which pit bottoms were visible (Jungblut *et al.* 2016). Thus, all three sample layers were likely exposed to at least some PAR. Oxygen gradients were measured in pits versus ridges (unpublished data), and the bottoms of pits were anoxic in contrast to the presence of 7.4 µmol O_2_/ L in the water column immediately above the mat (Jungblut *et al.* 2016). Thus, the mat spanned the local oxycline, although specific [O_2_] for each sample type is not well constrained. Even though PAR and [O_2_] are poorly constrained in detail, some illumination of all sample types is consistent with the presence of cyanobacteria, diatoms, and anoxygenic phototrophs in all layers. Furthermore, there are no statistically significant variations in photoautotrophs, specifically Cyanobacteria, among the mat layers (Figure 4). Rather, the OTUs that vary significantly between layers at 9.3 m are putative aerobes (physiology as reported in (Finster, Liesack and Tindall 1997; Schneiker *et al.* 2007). These aerobes are more abundant in the top samples, which were predicted to be exposed to more O_2_. Thus, [O_2_] likely structured the community composition at 9.3 m, with the communities constructing both mat topography and the local redox gradient.

The film, top, middle, and bottom mat layers at 9.8 m hosted similar communities to each other, although the film samples were dominated by *Phormidium pseudopristleyi*. The seasonal variability in [O_2_] at 9.8 m in Lake Fryxell, as well as the proximity of the oxycline, allow for the presence of sulfide (Jungblut *et al.* 2016). The dominance of *P. pseudopristleyi* as a primary producer at 9.8 m suggests that it is well adapted to the low light and anoxia in this habitat. Specifically, it appears to tolerate the presence of sulfide (Jungblut *et al.* 2016), unlike most cyanobacteria (Cohen *et al.* 1986; Miller and Bebout 2004). The lack of variation in other microbial guilds and the paucity of *P. pseudopristleyi* at other water depths (Figure 2) suggest that PAR structured the communities and that the small amount of seasonal O_2_ (Sumner *et al.* 2015) had little effect on the distribution of OTUs.

The effects of PAR on microbial mats are well documented, and many microbial mats are structured by redox cycles that depend on photosynthetic organisms (*e.g.*, Whitton 2012 and references therein). The microbial mats at 9.0 m and 9.8 m provide insights into the extent that PAR affects the composition and diversity of a microbial community in the absence of physiologically meaningful variations in [O_2_]. All layers at 9.0 m are supersaturated with oxygen, and PAR decreases from the top to bottom of the microbial mat. Similarly, layers at 9.8 m mostly lack oxygen, and PAR decreases from the top to bottom of the microbial mat. In both habitats, the decrease in the ratio of photoautotrophs to heterotrophs with depth into the mats suggests that PAR affects the microbial community by allowing photoautotrophs, specifically cyanobacteria, to dominate the illuminated top layers while heterotrophs, specifically proteobacteria, dominate the darker underlying layers. These results are therefore consistent with PAR structuring communities largely by influencing the distribution of photoautotrophs. This trend of cyanobacteria in top layers and proteobacteria in underlying layers is found in other microbial mat ecosystems (Kunin *et al.* 2008; Harris *et al.* 2013; Bernstein *et al.* 2017; Maier *et al.* 2018), raising the possibility that alpha diversity in microbial ecosystems is generally more dependent on the distribution of net primary producers than heterotrophs.

PAR further modulates microbial communities via the photosynthetic conversion of PAR into net primary productivity (NPP). Most organic carbon in MDV lakes is autochthonous (54), implying a direct relationship between NPP and PAR in Lake Fryxell. In these benthic mats, PAR and alpha diversity are negatively correlated (Figure 2), consistent with the hypothesis that microbial ecosystems in the photic zone function more efficiently with a less diverse community, one dominated by oxygenic photosynthesizers (*e.g.*, Hurlbert and Stegen 2014).

### Scale-dependent relationships between energy and diversity

We observed variable relationships between diversity, community composition, PAR and [O_2_] in Lake Fryxell. Diversity decreases with decreasing energy (PAR) input at the meter-scale across the lake floor in accordance with the species-energy theory, which states that diversity increases with greater energy capture by a community (Hurlbert and Stegen 2014). However, at the millimeter-scale of the layers, the dominance of photoautotrophs suppresses alpha diversity in two of the habitats (the 9.0 m top layer and the 9.8 m film and top layers), resulting in a negative correlation between PAR and diversity. Thus, across large-scale changes in PAR and [O_2_] in Lake Fryxell, species energy-theory explains community composition and diversity. However, at a smaller scale, increases in diversity into mat layers may follow the maximum power principle, by which community diversity and composition shift to optimize energy consumption under prevailing biogeochemical constraints (DeLong 2008). Specifically, photoautotrophs provide the bulk of the NPP, but their abundance reduces the diversity of the top mat layer; the absence of oxygen in deeper layers requires that anaerobic metabolisms to optimise energy use. The community structure of mats in Lake Fryxell therefore reflects the optimization of energy use across small scale-PAR and [O_2_] gradients as well as increased diversity in response to more energy availability across large-scale PAR and [O_2_] gradients.

Chase and Leibold (Chase and Leibold 2002) suggested a highly variable relationship between productivity and diversity that changes depending on how large a landscape is considered, a notion recently supported by studies of terrestrial plant communities (Grace *et al.* 2016). Similar to plant and animal communities, microbial mat communities in Lake Fryxell are structured in a scale-dependent manner. The differences in Lake Fryxell between meter-scale and millimeter-scale community diversity and composition support the hypothesis that integrating ecological theories at different spatial scales may best explain observed communities (Leibold and Chase 2017). Microbial mats therefore provide tractable systems within which to study the complex feedbacks among organisms and between communities and their environment.

In Lake Fryxell, the benthic microbial mats contain diverse taxa, and they respond to the spatially heterogeneous features of their habitats, specifically PAR and [O_2_], both in terms of phylogenetic diversity and relative population abundances. Alpha diversity is greatest where [O_2_] is highest and PAR is absent. Differences in population abundances are also strongly correlated to [O_2_], with greater beta diversity in oxygenated habitats and at low PAR. One OTU, *P. pseudopristelyi*, dominates the film layer at 9.8 m, which suggests that it is better suited to live in the presence of trace hydrogen sulfide near the oxycline than the majority of other Cyanobacterial genera living in other parts of the lake. These results, based in PCR-free sequencing, more biological replicates than previously recovered, and protein-level taxonomic characterization, support the patterns discovered in recent amplicon-based surveys (Jungblut *et al.* 2016).

In summary, where [O_2_] varies minimally in Lake Fryxell, either where it saturates the microbial mats (9.0 m) or where it is absent (middle and bottom layers at 9.8 m), PAR structures the overall microbial community from top to bottom. But where [O_2_] varies spatially, the microbial community is structured by [O_2_]. The main source of energy in Lake Fryxell is PAR, but [O_2_] influences phylogenetic diversity and taxonomic composition to a greater degree. The response of microbial mat communities to their geochemical environment is scale-dependent, as is the case in plant and animal communities, and microbial ecosystems can provide important test cases for ecological theories.

## Acknowledgements

Logistic field work support was provided by the National Science Foundation Division of Polar Programs through the McMurdo Long Term Ecological Research project (grants OPP-115245 and OPP-1637708) and Antarctic New Zealand Program. Field assistance was provided by Colin Hillmann. Support for genomics and data analysis was provided by NASA Astrobiology through grant NN13AI60G. The authors declare no competing financial interests in relation to the work described.

---

**Table S1.** Summary statistics of sequence data for samples

| Sample Type | n | Total bp | Total Reads | Average Length (bp) |
| --- | --- | --- | --- | --- |
| 9.0 Top | 16 | 4.19E+09 | 1.56E+07 | 268 |
| 9.0 Middle | 20 | 6.38E+09 | 2.39E+07 | 266 |
| 9.0 Bottom | 16 | 5.89E+09 | 2.23E+07 | 264 |
| 9.3 Top | 15 | 4.91E+09 | 1.85E+07 | 265 |
| 9.3 Middle | 15 | 5.43E+09 | 2.05E+07 | 264 |
| 9.3 Bottom | 16 | 6.56E+09 | 2.47E+07 | 266 |
| 9.8 Film | 5 | 2.09E+09 | 7.82E+06 | 268 |
| 9.8 Top | 13 | 5.50E+09 | 2.07E+07 | 266 |
| 9.8 Middle | 14 | 3.48E+09 | 1.31E+07 | 266 |
| 9.8 Bottom | 11 | 2.83E+09 | 1.07E+07 | 266 |

**Table S2.** Simpson’s diversity indices and species richness

| Sample Type | Simpson's Diversity Index | Species Richness |
| --- | --- | --- |
| 9.0 m Top | 0.616 | 17,008 |
| 9.0 m Middle | 0.7 | 18,181 |
| 9.0 m Bottom | 0.656 | 19,548 |
| 9.3 m Top | 0.655 | 17,383 |
| 9.3 m Middle | 0.698 | 18,044 |
| 9.3 m Bottom | 0.701 | 18,295 |
| 9.8 m Film | 0.715 | 14,561 |
| 9.8 m Top | 0.759 | 17,590 |
| 9.8 m Middle | 0.731 | 17,213 |
| 9.8 m Bottom | 0.707 | 16,609 |

**Table S3.**
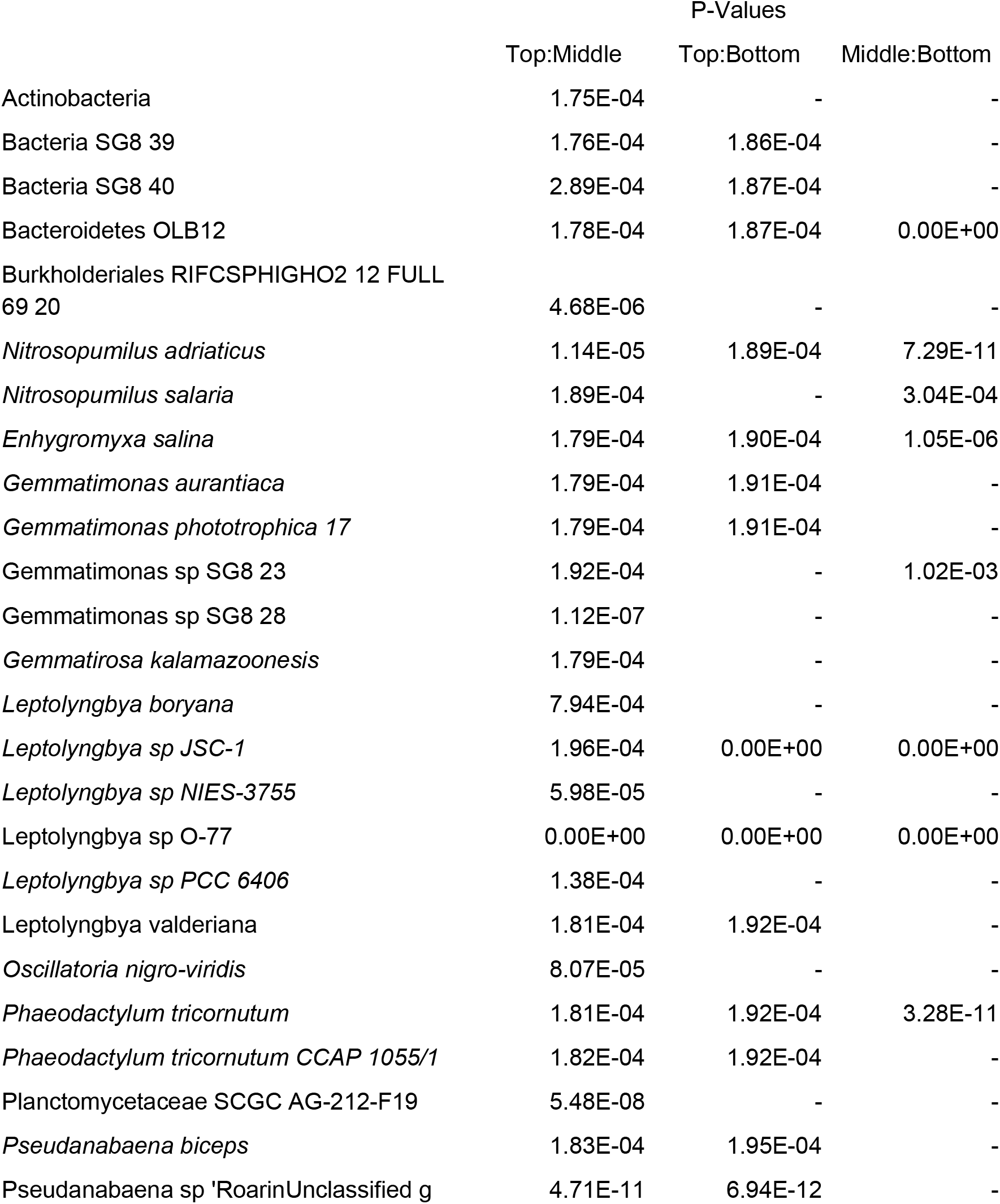
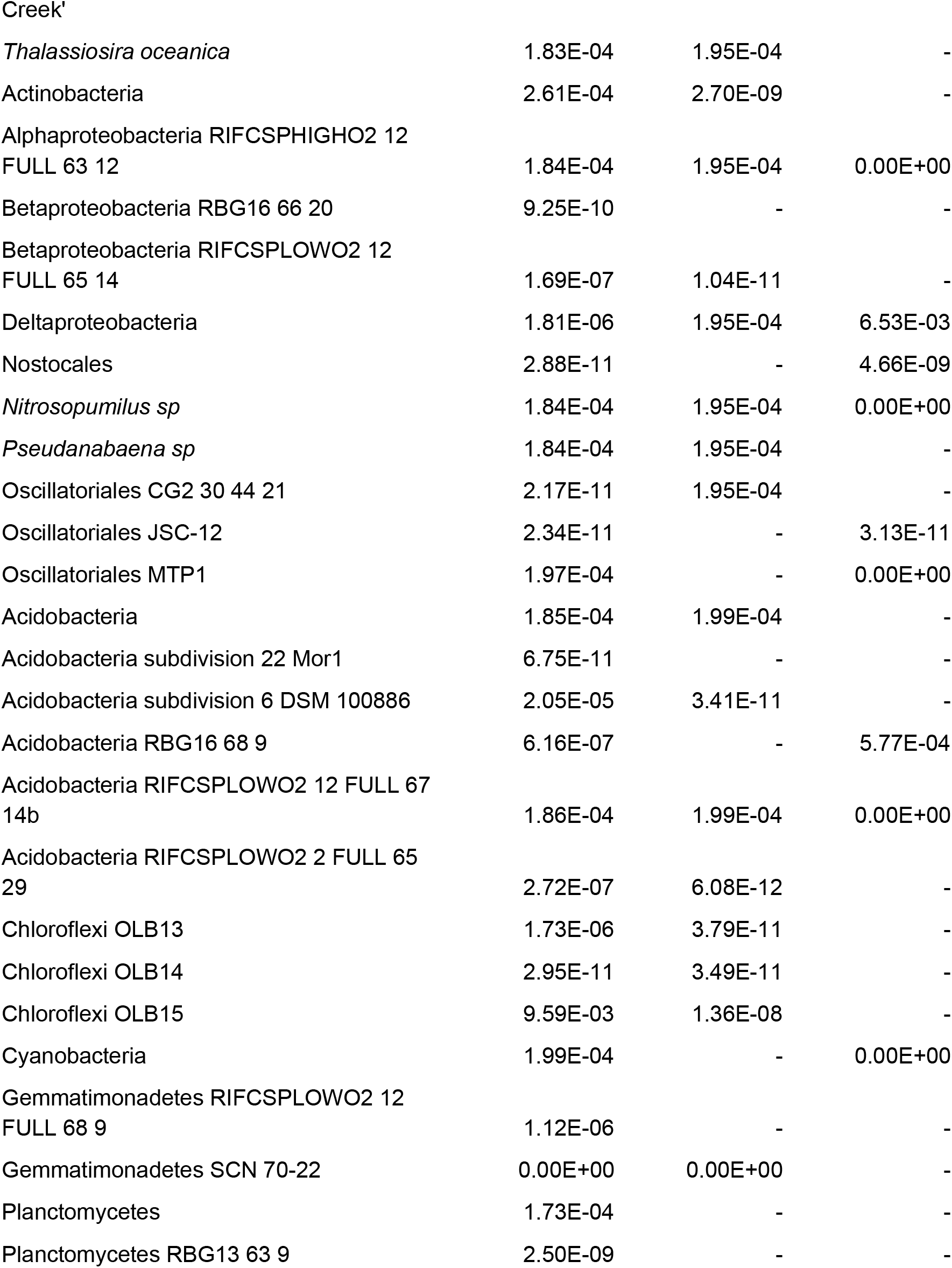
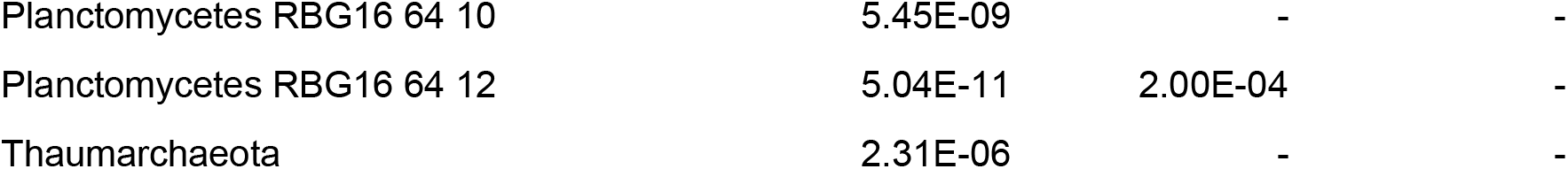
Taxa that differ significantly (P-value < 0.01) in relative abundance between layers at 9.0 m. Adjusted P-values were calculated using Tukey’s HSD. No difference is marked with -.

**Table S4.**
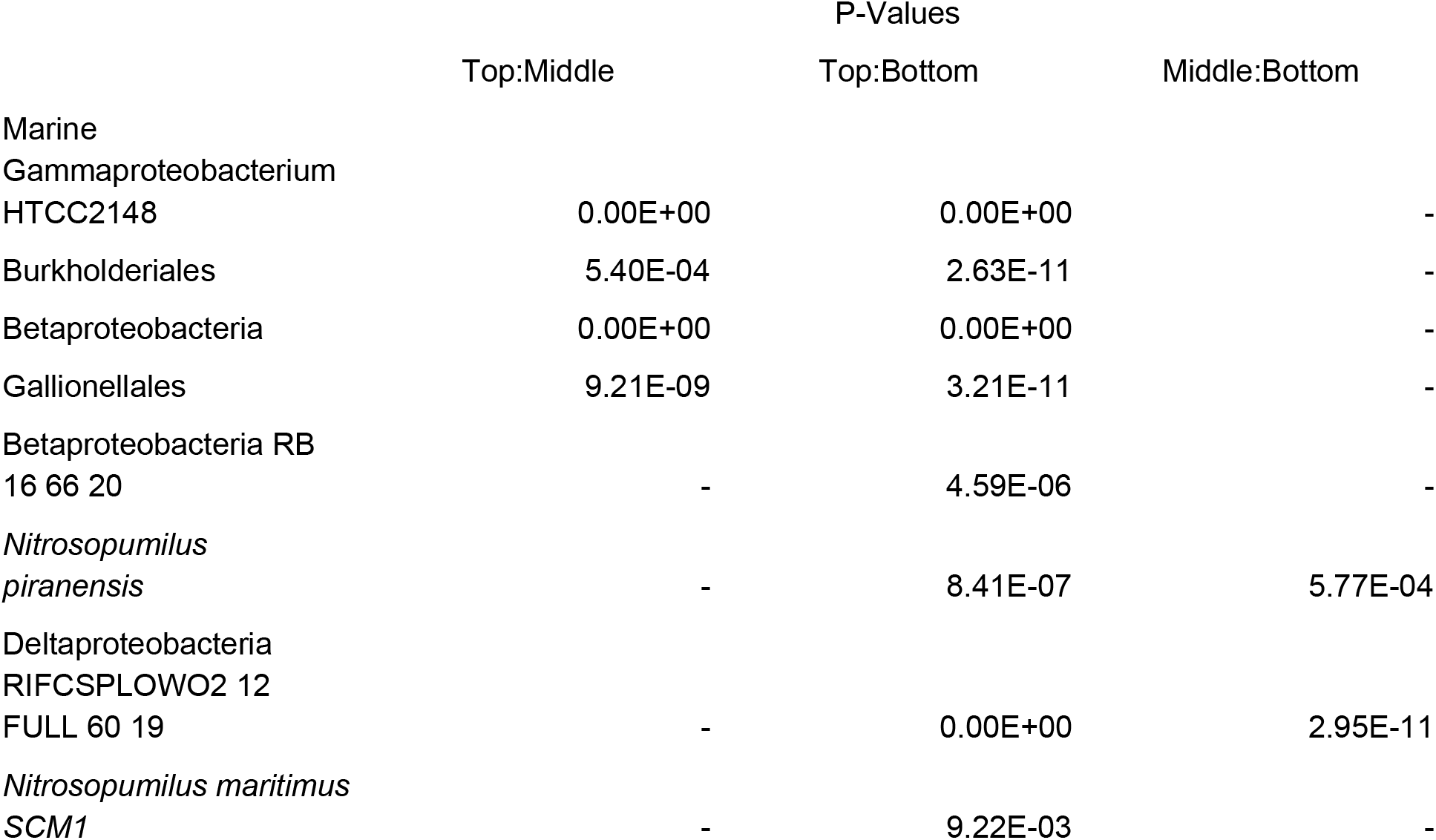
Taxa that differed significantly (P-value < 0.01) in relative abundance between layers at 9.3 m. Adjusted P-values were calculated using Tukey’s HSD. No difference is marked with -.

**Table S5.** Taxa that differed significantly (P-value < 0.01) in relative abundance between layers at 9.8 m. Adjusted P-values were calculated using Tukey’s HSD.

|  | P-Values |  |  |  |  |  |
| --- | --- | --- | --- | --- | --- | --- |
|  | Film:Top | Film:Middle | Film:Bottom | Top:Middle | Top:Bottom | Middle:Bottom |
| *Phormidium*<br>*pseudopristleyi* | 6.02E-13 | 6.02E-13 | 6.02E-13 | 6.02E-13 | 6.02E-13 | 0 |

